# A new transcriptional metastatic signature predicts survival in clear cell renal cell carcinoma

**DOI:** 10.1101/2022.06.28.497333

**Authors:** Adele M. Alchahin, Shenglin Mei, Ioanna Tsea, Taghreed Hirz, Youmna Kfoury, Douglas Dahl, Chin-Lee Wu, Alexander O. Subtelny, Shulin Wu, David T. Scadden, John H. Shin, Philip J. Saylor, David B. Sykes, Peter V. Kharchenko, Ninib Baryawno

## Abstract

Clear cell renal cell carcinoma (ccRCC) is the most common type of kidney cancer in adults. When ccRCC is localized to the kidney, surgical resection or ablation of the tumor is often curative. However, in the metastatic setting, ccRCC remains a highly lethal disease. Here we take advantage of fresh patient samples that include treatment-naive primary tumor tissue, matched adjacent normal kidney tissue, as well as tumor samples collected from patients with bone metastases. Single-cell transcriptomic analysis of tumor cells from the primary tumors revealed a distinct transcriptional signature that was predictive of metastatic potential and patient survival. Analysis of supporting stromal cells within the tumor environment demonstrated vascular remodeling within the endothelial cells and a proliferative signature within the fibroblasts that was associated with poor survival. An *in silico* cell-to-cell interaction analysis highlighted the CXCL9/CXCL10-CXCR3 axis and the CD70-CD27 axis as potential therapeutic targets. Our findings provide biological insights into the interplay between tumor cells and the ccRCC microenvironment.

## INTRODUCTION

Renal cell carcinoma is the most common renal tumor in adults, and the clear cell subtype (ccRCC) accounts for 75-85% of all cases^1^. While localized disease can often be cured with surgical resection or thermal ablation, there is evidence of metastatic disease in approximately one quarter of patients at the time of diagnosis^2^. Tumor cells that arise initially as a clonal expansion of transformed cells then propagate in the ecosystem of the tumor microenvironment (TME) where distinct cell populations engage in complex interactions that promote tumor growth and metastatic spread^3^.

The TME is composed of stromal cells including immune cells, endothelial cells, fibroblasts, smooth muscle cells and pericytes^4^. Biological processes within the TME such as inflammation, hypoxia, angiogenesis, and epithelial-to-mesenchymal transition (EMT) contribute to tumor complexity and evolution, and may play a critical role in promoting distant metastasis^5^. Specific components of the TME are well established as therapeutic targets. One treatment strategy in ccRCC is to inhibit the VEGF signaling that is commonly dysregulated following VHL gene inactivation within RCC tumor cells and would otherwise stimulate endothelial cell growth and angiogenesis^6^. Immune checkpoint blockade (ICB) targets T-cells within the immune TME and has improved outcomes in patients with ccRCC both as monotherapy and in combination with other agents^6-8^. However, resistance to treatment is common and may partly be attributed to other tumor-protective roles of the microenvironment^7^. Improving therapies for metastatic RCC will require a deep understanding of how the tumor cells are specifically interacting with their microenvironment.

Comprehensive genomic studies in ccRCC have provided insights into the somatic alterations that affect tumor progression^9,10^ and response to immune checkpoint blockade^11^. Single-cell RNA-seq (scRNA-seq) studies on human RCC have provided immune cell atlases based on tumor samples from treatment naïve and treated patients. This has improved our understanding of the cell composition and cellular states of tumor-infiltrating immune cells that may contribute to the response to immunotherapy^12,13^. These studies highlight the proportion of exhausted CD8+ T cells and the function of immunosuppressive M2-like macrophages in advanced RCC^12^. While an infiltration of cytotoxic CD8+ T (CTLs) cells has been associated with an improved prognosis other solid tumor type, it has been correlated with a worse prognosis in ccRCC^14^. ICB treatment in RCC can remodel the microenvironment and can modify the interplay between cancer cells and immune populations such as CD8+ T cells and macrophages but patient responses to immune checkpoint blockade are still far from universal^15-22^. These studies have expanded our understanding of the immune components of the microenvironment though the composition of stromal cell populations and their interactions with tumor cells remain unclear. Elucidating the specific relationship between stromal cells and tumor cells within the TME will advance our understanding of carcinogenesis and cancer progression.

In this study, we profiled human ccRCC tumors and their matched normal control kidney tissue from treatment-naïve patients, in addition to primary tumors from patients presented with bone metastases at diagnosis. This single-cell transcriptomic analysis led to the following observations: (1) tumor cells are transcriptionally similar to a subset of proximal tubule cells which may be an indication of the tumor cell of origin^23^ (2) the synchronous comparison of primary tumor and bone-metastatic tumor tissues from two patients who presented with *de novo* metastases revealed a specific metastatic signature associated with poor prognosis, (3) the stromal cells within RCC tumors show the highest transcriptional difference of analyzed cell types when compared to adjacent normal kidney, and (4) building on other scRNA-seq studies of human RCC^12,13,23,24^, we identify additional components of the TME including a cellular map of the stromal cell compartment, and of cell-to-cell interactions within the tumor that might be vulnerable to therapeutic targeting. This careful dissection of the cellular and molecular landscape of ccRCC is intended to facilitate new avenues of therapeutic intervention and ultimately better treatments for patients suffering from ccRCC.

## RESULTS

### Primary human ccRCC show consistent microenvironmental changes as compared to matched normal kidney tissue

To provide an overview of the molecular and cellular landscape of patient-matched normal kidney and ccRCC tissue, we performed scRNA-seq profiling (10x Chromium) from freshly resected primary ccRCC tumors (16) and adjacent normal samples from 10 patients (Fig. 1a). Nine patients were diagnosed with ccRCC and 1 patient with papillary RCC, pRCC (pRCC was excluded from analysis). Two patients (RCC-BM1-PT and RCC-BM2-PT-1,2) had clinical metastases at multiple sites at the time of diagnosis (Extended Data Fig. 1a, Supplementary Table 1). Tumor tissues and adjacent normal kidney tissue collected from the same patient (from two patients, tumor in different location on the kidney were profiled) permitted a matched comparison and helped to control for inter-individual variation. In total, we sequenced 157,881 cells (ccRCC: 122,054 cells + normal kidney: 35,827 cells) and samples were integrated using a joint analysis of the heterogenous samples (Fig. 1b and Extended Data Fig. 1b).

**Figure 1:**
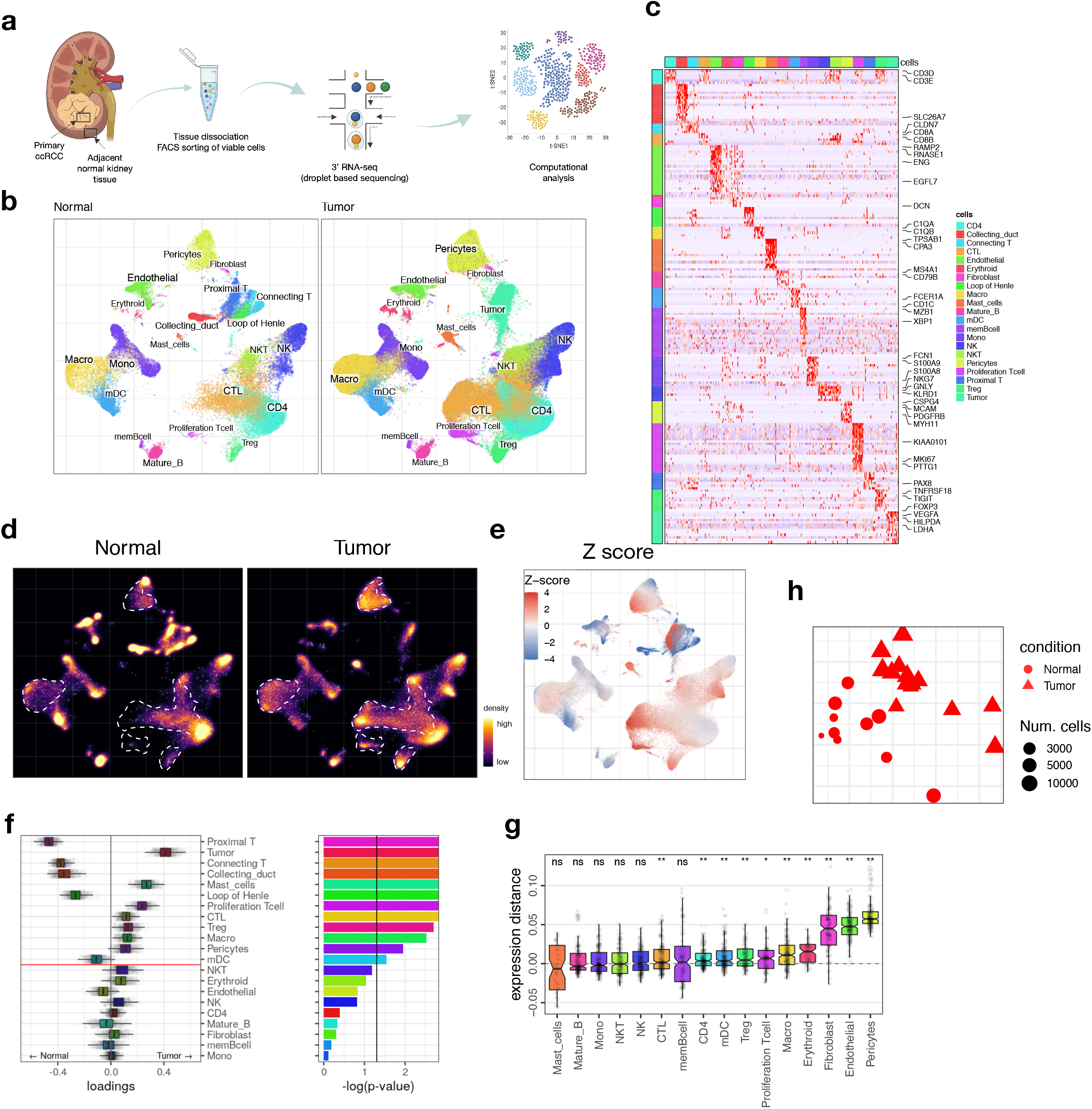
Single-cell landscape of the ecosystem in primary ccRCC and adjacent normal tissue. **a –** Experimental design. **b** – Integrative analysis of scRNA-seq samples from 11 RCC patients, visualized using a common UMAP embedding for adj-normal (left) and tumor kidney tissue (right). **c** – Heatmap showing expression of markers for major cell populations. **d** – Changes in the composition of all compartments combining all sample fractions and is visualized as cell density on the joint embedding. **e**- Statistical assessment of the cell density differences comparing tumor with adjacent normal. Wilcoxon test was used, visualized as a Z score. Red indicates increased cell abundance in tumor, blue indicates decreased cell abundance in tumor. **f** – Change in cell composition evaluated by Compositional Data Analysis. The x axis indicates the separating coefficient for each cell type, with the positive values corresponding to increased abundance in tumor, and negative to decreased abundance. The boxplots and individual data points show uncertainty based on bootstrap resampling of samples and cells. **g**– The boxplots showing the magnitude of transcriptional change between primary RCC and normal kidney tissue in major cell populations. The magnitude is assessed based on a Pearson linear correlation coefficient, normalized by the medium variation within primary RCC and normal kidney tissue. Statistics significance within each cell type is measured with permutation in sample group (see Methods). Boxplots: center line: median; box limits: upper and lower quartiles; whiskers extend at most 1.5x interquartile range past upper and lower quartiles. **h** – A t-SNE embedding of different samples, based on their overall expression distance. The similarity measure measures the magnitude of expression change for each subpopulation, using size-weighted average to combine them into an overall expression distance that controls the compositional differences. Shape indicates different sample fractions.

Unsupervised clustering identified 21 distinct clusters including normal kidney cell populations, tumor cells, immune and non-immune stromal populations (Fig. 1b). The stromal cells included pericytes (expressing *RGS5, MYH11*), endothelial cells (*RAMP2, CD34*) and fibroblasts (*DCN, LUM*). Lymphoid cells included T cells (*CD3D, CD3E*), NK cells (*KLRD1, XCL1*), and B cells (*CD79, CD19*). The myeloid compartment consisted of macrophages (*CD68, C1QA, C1QB, C1QC*), monocytes (*FCN1, S100A9*) and myeloid dendritic cells (*CLEC9A, CD1C*) (Fig. 1c).

The patient-matched adjacent kidney samples (normal, tumor uninvolved) allowed us to identify tumor specific changes. Cell fraction differences were performed using cell density analysis on the joint UMAP embedding and direct comparison of cell proportions. At the global level, this analysis revealed an enrichment of pericytes in the RCC as compared to their adjacent normal kidney tissues, an increase in the CTLs and proliferating T cells, as well as macrophages (Fig. 1d-e and Extended Data Fig. 1c-d). As proportional changes of one subtype could potentially skew the representation of other subtypes, we confirmed findings via a Compositional Data Analysis technique to estimate compositional changes^25^ (Fig 1f, Extended data Fig. 1d).

In addition to the changes in the proportion of cell populations, we examined transcriptional state differences between tumor and adjacent normal kidney tissues using an expression distance measurement based on the Pearson linear correlation. The stromal compartment, including fibroblasts, endothelial cells and pericytes, demonstrated the largest transcriptional differences between tumor and adjacent normal (Fig. 1g). Specifically, there was upregulation of genes associated with cell motility and angiogenesis (blood vessel morphogenesis) (Extended data Fig. 1e-f). This suggests cancer-specific alterations in the stromal microenvironment during tumor progression. Expression distances were further projected using multidimensional scaling (Fig. 1h), resulting in consistent separation of normal kidney tissue (circular) from primary RCC (triangular).

### ccRCC tumors establish an immunosuppressive tumor microenvironment

Several types of cancer, including RCC, are heavily infiltrated by immune cells even when localized ^12,13^. Subcluster analysis of myeloid cells identified two subpopulations of myeloid dendritic cells (mDC), three populations of monocytes (Mono-1, 2, 3), and three populations of macrophages (Macro-1, 2, 3) (Fig. 2A). In the mDCs group, CD1C^+^ mDC (*CD1C, FCER1A, CLEC10A*) (Extended Data Fig. 2a) were reduced in the tumor compartment compared to the adjacent kidney (Fig. 2b.

**Figure 2:**
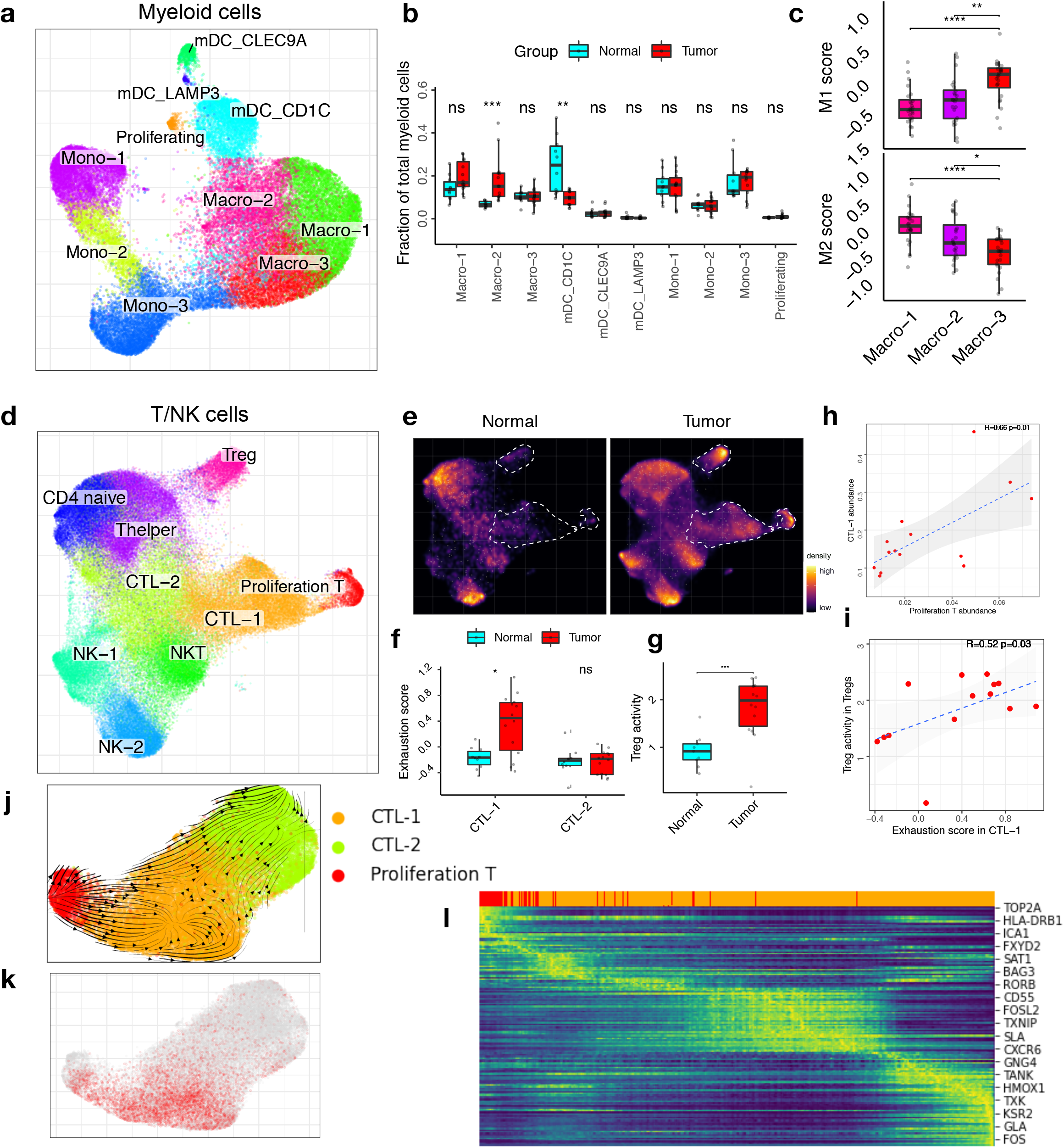
An immunosuppressive environment in ccRCC. **a**- UMAP embedding demonstrating myeloid subpopulations. **b**- Boxplots showing the proportions of myeloid subsets divided by the total myeloid cell number (*p<0.05, ***p<0.001) for select pairs, based on a Wilcoxon rank sum test. **c**- Average expression of the M1 and M2 macrophage signature gene panel across different monocyte populations shown as boxplot. Statistics are accessed using Wilcoxon rank sum test (*p<0.05, ****p<0.0001). **d-** UMAP embedding showing T cell subpopulations. **e**- Changes in the composition of the myeloid compartment between tumor and normal is visualized as cell density on the joint embedding. **f**- Boxplot presenting the exhaustion score of the T cell population comparing the adjacent normal kidney samples (turquoise) with tumor samples (red). Statistics are accessed using Wilcoxon rank sum test. **g**- Boxplots illustrate significant increase of Treg activity in the primary RCC. Boxplots in (b-c) and (f-g) include centre line, median; box limits, upper and lower quartiles; and whiskers are highest and lowest values no greater than 1.5x interquartile range. **h**- Correlation of proliferation T cells abundance and CTL-1 abundance is shown as scatter plot. Pearson linear correlation estimate and p-values are shown. **i**- Correlation of exhaustion signature score in CTL-1 and Treg activity score in Tregs is shown as scatter plot. Pearson linear correlation estimate and p-values are shown. **j**- RNA velocity analysis of the transitions of CTL-1, CTL-2 and proliferating T cells. **k**- Visualization of T cell exhaustion score shown on UMAP embedding **l**- Expression trends of the top 200 genes whose expression correlates with velocity pseudotime in j.

We identified an increasing population of proliferating myeloid cluster expressing both mDC and macrophage signatures (*MKI67, KIAA0101, CD1C, C1QA*) (Extended data Fig. 2c-d). Focused sub-cluster analysis of these cells revealed two clusters: proliferating macrophage (*C1QA, APOC1*) and proliferating mDC (*CLEC10A, CD1C*); the proliferating macrophages showed a similar cell phenotype to Macro-1/2 expressing *CD163, TREM2* and *SPP1* (Extended data Fig. 2a-b).

The three macrophage subpopulations showed a distinct gene signature (Macro-1: *SEPP1, PDK4, FCGR1A*; Macro-2: *SPP1, CXCL9, CXCL10, CD68*; Macro-3: *PLAUR, IL1B, CXCL2*) (Extended data Fig. 2b). Macro-1 and Macro-2 showed a significantly increased M2 macrophage signature score compared to Macro-3 (Fig. 2c) and expressed typical M2 marker genes (*CD68, TGFB1, CD163*^11,26-28^, *TGFB2*^29^, *CCL18*^30^, *MMP14*^31^, *CTSD*^32^, *MARCO*^*33*^ and *CSF1R*^34^) (Extended data Fig. 2a) suggesting that these macrophages are suppressive of the immune response and likely supporting tumor growth^13^. Macro-3 showed a significantly higher M1 signature score compared to Macro-1 and Macro-2 (Fig. 2c) and expressed high levels of *IL1A* and *IL1B*^12^ (Extended data Fig. 2a) indicative of a proinflammatory state. Macro-2 was separated from the other macrophage clusters by over-expressing *TREM2* and *SPP1* (Extended data Fig. 2a-b), two genes that have been associated with tumor angiogenesis and immune checkpoint therapy^35^. The expression of *SPP1*^*+*^ tumor associated macrophages has previously been identified in eight other tumor types, including colorectal cancer and breast cancer^35^. The overexpression of *TREM2*^*28*^ in macrophages in tumors has been linked to resistance to immune checkpoint therapy^36^. Consistent with this, *SPP1* and *TREM2* expression were significantly increased in the tumor fraction compared to the adjacent normal kidney tissue (Extended data Fig. 2e). TREM2 is exclusively expressed in the macrophage population (Extended data. 2f), and further analysis of bulk RNA-seq data^37^ shows that *TREM2-high* tumors are associated with poor survival outcomes (Extended data Fig. 2j-k), suggesting that TREM2+ M2 macrophages play an important role in ccRCC progression. Furthermore, we validated the increased TREM2+ macrophage in tumor using flow cytometric anlaysis (Extended data Fig. 2g, i), and found infiltrated TREM2+ cells within the ccRCC tumor microenvironment by utilizing the public ccRCC spatial transcriptomic data^38^ (Extended data Fig. 2h).

Subcluster analysis of the T lymphocytes revealed the anticipated T cell subpopulations including CD8^+^ cytotoxic T lymphocytes (CTLs) (*CD8A, IFNG*), T_regs_ (*IL2RA, CTLA4, FOXP3*), CD4^+^ naïve T cells (*CCR7*), T helper cells (Th) (*RORC, IL17A*), and subgroups of NK cells (*NKG7, NCR1*) (Fig. 2d; Extended data Fig. 3a). We observed proliferating T cells (*MKI67, CD8A, TOP2A*), and two different CTL populations: CTL-1 (*HAVCR2, PDCD1*) and CTL-2 (*CD8A, KLRG1*) (Fig. 2d; Extended data Fig. 3a). In comparison to the adjacent kidney, the proportion of CTL-1 and proliferating T cells were significantly increased in the tumor fraction (Fig. 2e; Extended data Fig. 3b)^12^. Further, CTL-1 expressed known immune-inhibitory molecules such as *PDCD1, TOX, HAVCR2, LAG3* and *CTLA4* indicating that the CTL-1 cells are suppressed in RCC^12^ (Fig. 2f; Extended data Fig. 3a-d). The CTL-1 exhaustion score^39^ was significantly higher in tumor tissue compared to adjacent kidney (Fig. 2f), suggesting that the tumor-associated CTL-1 might have diminished function. In parallel, we observed an increased T_reg_ activity signature score in the tumor fraction (Fig. 2g), with association to CTL-1 exhaustion (Fig.2i; Extended data 3f; Extended data 9b).

**Figure 3:**
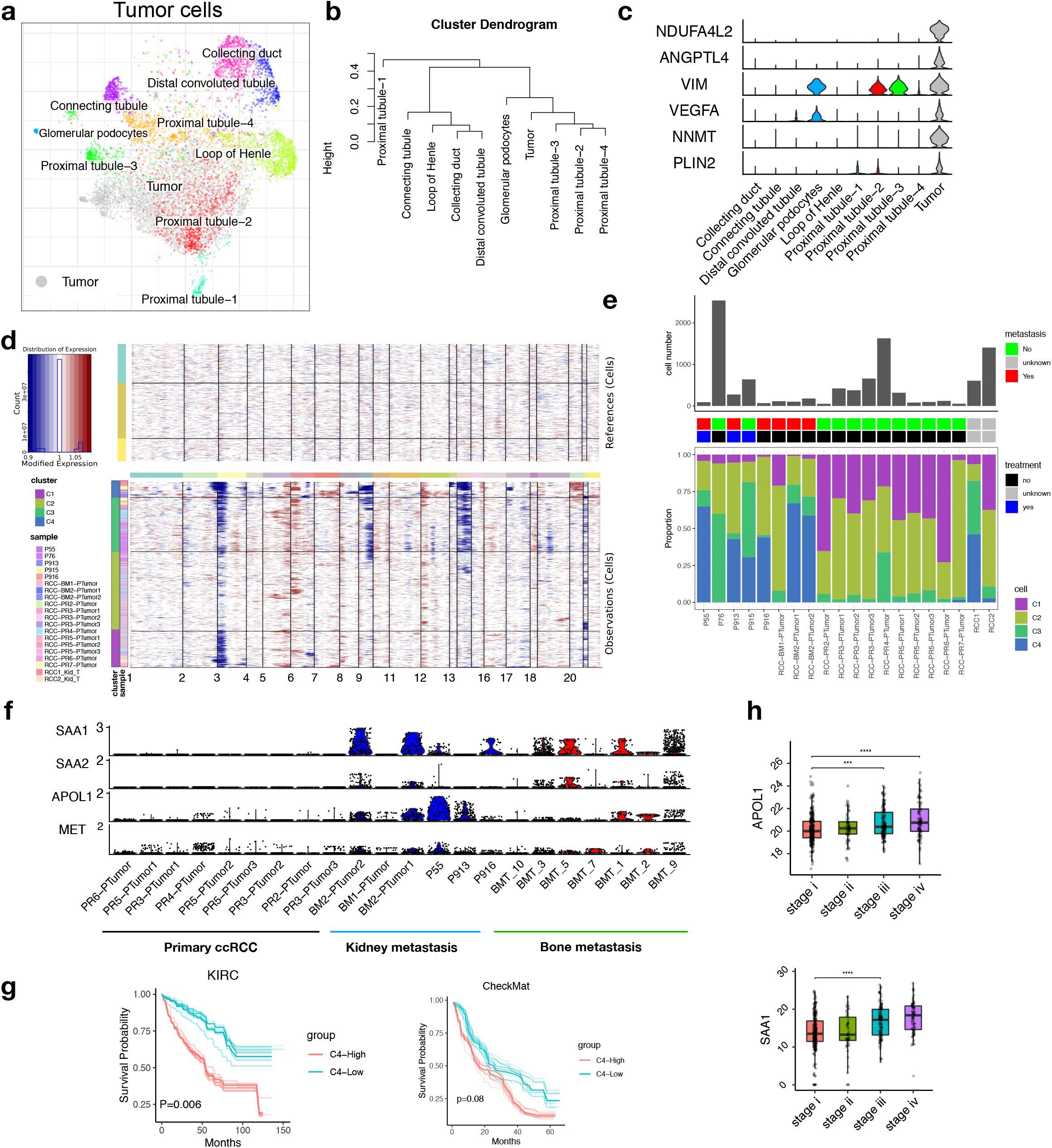
Intra-tumoral heterogeneity reveals distinctive malignant cell subclones. **a**- Joint alignment of nephron anatomy cells from normal kidney tissue and tumor cells from primary RCC tissue in UMAP embedding, colored by cell annotation. **b**- Expression distance of different cell subpopulations is shown as a dendrogram. **c**- Violin plot showing representative marker gene expression of tumor cells. **d**- Inferred CNV profile of tumor cells with normal nephron anatomy cells as normal reference. **e**- Summary of tumor cell subclones, number of tumor cells (Top), clinic pathological features (middle), cell abundance of tumor sub-clones (bottom) in each sample. **f**- Violin plot showing metastatic signature gene expression in patient tumor cells from primary ccRCC, local kidney metastasis and bone metastasis. **g**. Overall survival (OS) analysis for TCGA KIRC and CheckMate bulk RNA-seq data. Patients were stratified into two groups based on the average expression (binary: top 25% versus bottom 25%) of metastatic signatures as annotated by key marker genes in Figure 4f. Bootstrap resampling were performed on signature genes and p-value was calculated using the 95% reproducibility power p-value (Methods). **h-** Expression of APOL1 and SAA1 are shown as boxplot, stratifying patients by disease stage (TCGA KIRC) with Wilcoxon rank sum test (****p<0.0001). Boxplots: center line: median; box limits: upper and lower quartiles; whiskers extend at most 1.5x interquartile range past upper and lower quartiles.

Within the tumor fraction, CTL-1 abundance was significantly correlated with the proliferating T cell cluster (Fig. 2h). RNA velocity analysis can be used to infer precursor progeny cell dynamics^40,41^ and we identified a directional flow suggesting that the proliferating T cells give rise to the CTL-1 population (Fig. 2j-l; Extended data Fig. 3e).

The presence of NK cells in RCC may represent a critical component of the anti-tumor response and NK cell infiltration in patient tumors has been associated with an improved clinical prognosis^42^. Comparing the adjacent normal kidney tissue and the tumor, we detected two subpopulations: NK-1 and NK-2 (Fig. 2d) which were further annotated as CD56^dim^ NK-1 cells (*CD44, XCL1, XCL2, KLRC1*) and CD56^bright^ NK-2 cells (*FGFBP2, CX3CR1, GZMB*)^43^ (Extended data Fig. 3a, h-i). Using a cytotoxicity score to confirm that they were functionally distinct (Supplementary Table 2), we demonstrated that the CD56^bright^ subset of NK cells have a significantly greater cytotoxic phenotype (Extended data 3i) with expression of cytotoxic genes (*PRF1, GZMA, GZMB, GZMH, GZMM*)^43^ (Extended data Fig. 3b; h-i).

### A distinct metastatic tumor cell cluster correlates with poor prognosis

It is believed that primary ccRCC develops in the proximal part of the nephron^44^, and we focused on the adjacent normal kidney tissue where we identified four distinct sub-populations in the nephron proximal tubule (PT), PT1-4 (Extended data Fig. 4a-b). The PT1 transcriptional signature was characterized by metabolism-associated genes (*FABP1, PRODH2*)^45^ and the PT4 signature of kidney fibrosis (*MMP7*^46^ and *ITGB6*^47^) (Extended data Fig. 4b). Interestingly, in PT2 the transcriptional signature was one of inflammation and regeneration gene expression (*SOX9, IL32, VCAM1*) (Extended data Fig. 4b), which has been related to carcinogenesis and cancer progression^48,49^. Neighboring the proximal tubule cells was a small subpopulation of glomerular podocytes (*SEMA3G, CLDN5, CR1*)^50^ found in the Bowman’s capsule where they wrap around capillaries of the glomerulus (Extended data Fig. 4a-b).

**Figure 4:**
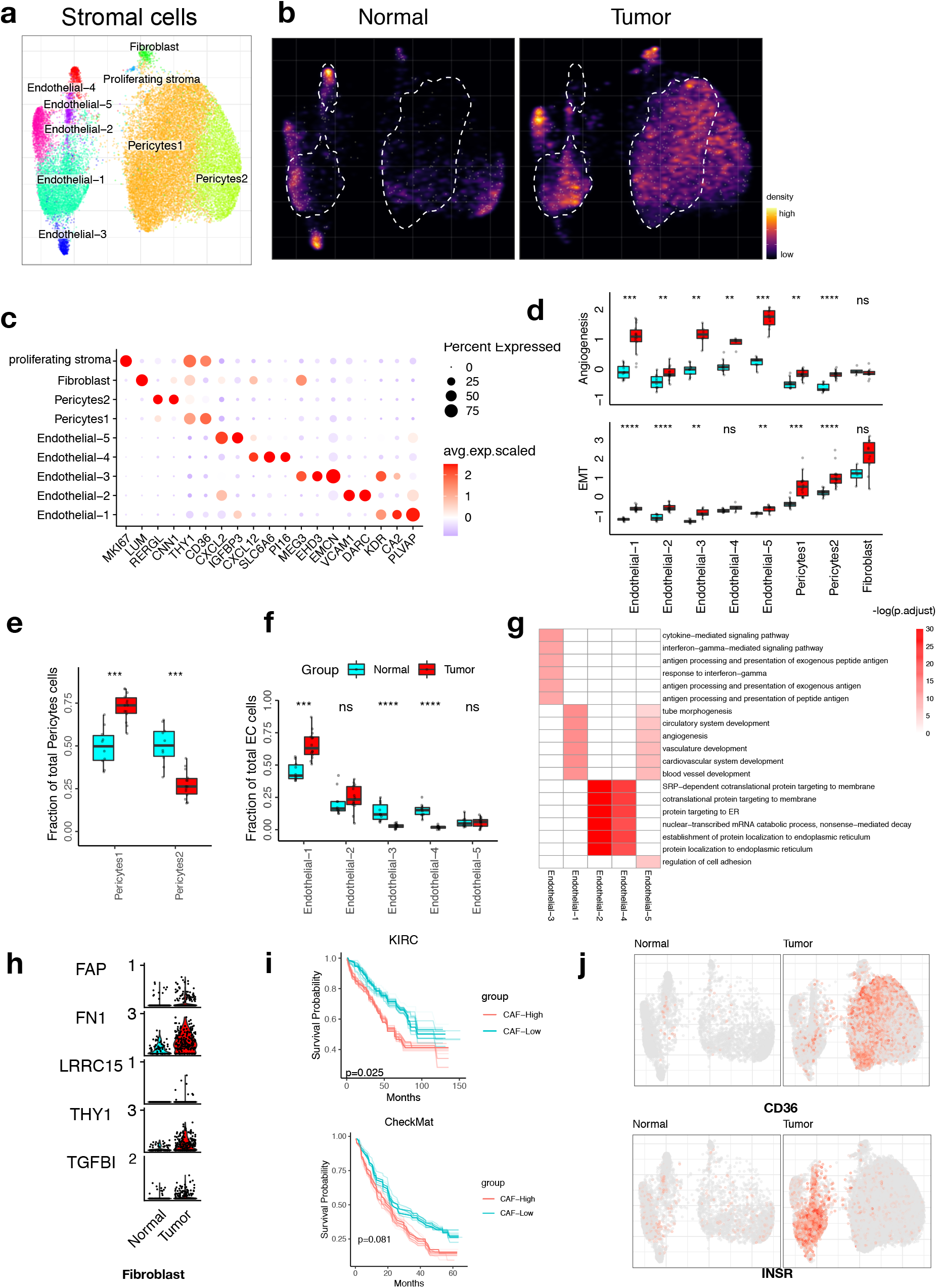
ccRCC is driven by vascular remodelling through angiogenic and EMT switch in stromal cells. **a**- UMAP embedding of stromal cells, color-coded by the cell subtypes. **b**- Changes in the composition of stromal cell is visualized as cell density on UMAP embedding. **c**- Dot plots showing representative marker gene expression across different stromal subsets. The color represents scaled average expression of marker genes in each cell type, and the size indicates the proportion of cells expressing marker genes. **d**- Boxplot plot illustrate EMT and angiogenesis signature score across different stromal cell subpopulations of normal kidney tissue and primary RCC. **e**- Boxplots showing the proportions of pericytes subsets divided by the total pericytes cell number. **f**- Boxplots showing the proportions of endothelial subsets divided by the total endothelial cell number. Boxplots in (d-f) include centreline, median; box limits, upper and lower quartiles; and whiskers are highest and lowest values no greater than 1.5x interquartile range. Statistics significance are accessed using Wilcoxon rank sum test (*p<0.05, ****p<0.0001). **g -** The enriched GO BP terms of top 200 upregulated genes for each stromal cell subtype comparing to adjacent normal. **h**- Violin plots showing cancer associated fibroblast (CAF) signature gene expression in tumor and adjacent normal fibroblast. **i**- Similar with Figure 3g, showing ccRCC samples with higher CAF signature gene expression have worse overall survival in TCGA KIRC (top) and CheckMate (bottom) data. **j-** INSR and CD36 expression in UMAP embedding separately for tumor and adjacent normal tissue.

To relate the malignant cells to normal kidney anatomy, we compared the transcriptional states of tumor cells and normal nephron cell subsets from adjacent normal kidney. Tumor cells and proximal tube cells showed a similar transcriptional profile and the malignant population clustered most closely with PT2 in the joint integration (Fig. 3a, b), pointing to PT2 as a possible tumor cell of origin. Young et al.^23^ found a subset of PT cells (VCAM1+, SLC17A3+ and SLC7A13-) could be the origin of ccRCC, aligned with our annotated proximal tubule 2 (PT2) and PT1 (Extended data Fig. 4c). We further utilize bulk RNA-seq data and found a significant upregulation of SOX9, IL32 and SLC22A6 (PT2), and downregulation of *SLC22A6/ SLC22A8 (*PT1) in tumor compared to adjacent normal tissue (Extended data Fig. 4d), confirming our cell origin hypothesis on PT2 cells.

Comparing the normal PT cells, we investigated differentially expressed genes with a focus on transcriptional programs known to underly tumorigenesis in ccRCC. Upregulated genes included *VEGFA, NDUFA4L2, NNMT2*, and *PLIN2*, all previously shown to be involved in tumor development and progression (Fig. 3c, Extended data Fig. 4c-d)^51,52^.

We next inferred large-scale chromosomal copy number variations (CNVs) based on transcriptomic data using inferCNV^53^. Our findings are in line with previous descriptions of frequent aberrations in ccRCC including the loss on chr3p, chr14 and gains on 5q (Fig. 3d; Extended data 5a)^9,54^. One papillary RCC sample (PR6) displayed a markedly different CNV profile (gain of chr3p, chr7, chr17) and was therefore removed from subsequent analysis.

Recent studies have revealed recurrent CNVs in patients with kidney cancer (TRACERx Renal and TCGA KIRC cohorts)^10,55^. To analyze tumor cell heterogeneity, we performed hierarchical clustering of inferred CNV profiles with a comparison to publicly available single-cell resolution datasets from advanced ccRCC patients^13,23^. Clustering of CNV profiles revealed four major tumor clusters that were shared by different patients (Fig. 3d; Extended data 5c). C1 and C2 was found mainly in primary RCC patients without metastatic disease, defined by loss of chromosome 3p (Fig. 3d-e), driven by genes associated with catabolic and metabolic processes (Extended data Fig. 5c). C3 was characterized by losses on chr3p, chr6p, chr14, and showed high expression of *APOE, APOC1, ACSM2A* and *ACSM2B* (Extended data 5d). C4 had the most abundant copy number aberrations and was notably enriched in the patients with multiple metastatic sites of disease (Fig. 3d-e). C4 also had a distinct transcriptional profile with upregulation of *SAA2, SAA1, APOL1, MET* (Extended data 5d), exclusive when compared to other cell types (Extended data Fig. 5e). *MET* is a tyrosine kinase receptor involved in cell proliferation, survival, and migration^56^; it is among the therapeutic targets of cabozantinib^57^. We further perform DE gene analysis and defined a metastatic signature gene set (*SAA1, SAA2, APOL1, MET*, **Methods**). Notably, we observed that the metastatic signature was significantly associated with poor prognosis in two independent RCC cohorts (Fig. 3g). Even the gene expression of the individual genes APOL1 and SAA1 (Fig. 3f, h) correlated with disease stage. In addition, we also validate the metastatic signature gene expression in tumor cells from bone metastasis sites using an independent study of the bone marrow renal cell carcinoma (RCC) metastasis (GSE202813). Our analysis of 7 RCC metastasis cases shows the upregulation of metastatic signature compared to primary tumor cells, further demonstrating the reliability of tumor metastatic signature. Taken together, we identified a distinct tumor cell cluster with a transcriptomic program associated with metastatic potential and poor survival^58^.

**Figure 5:**
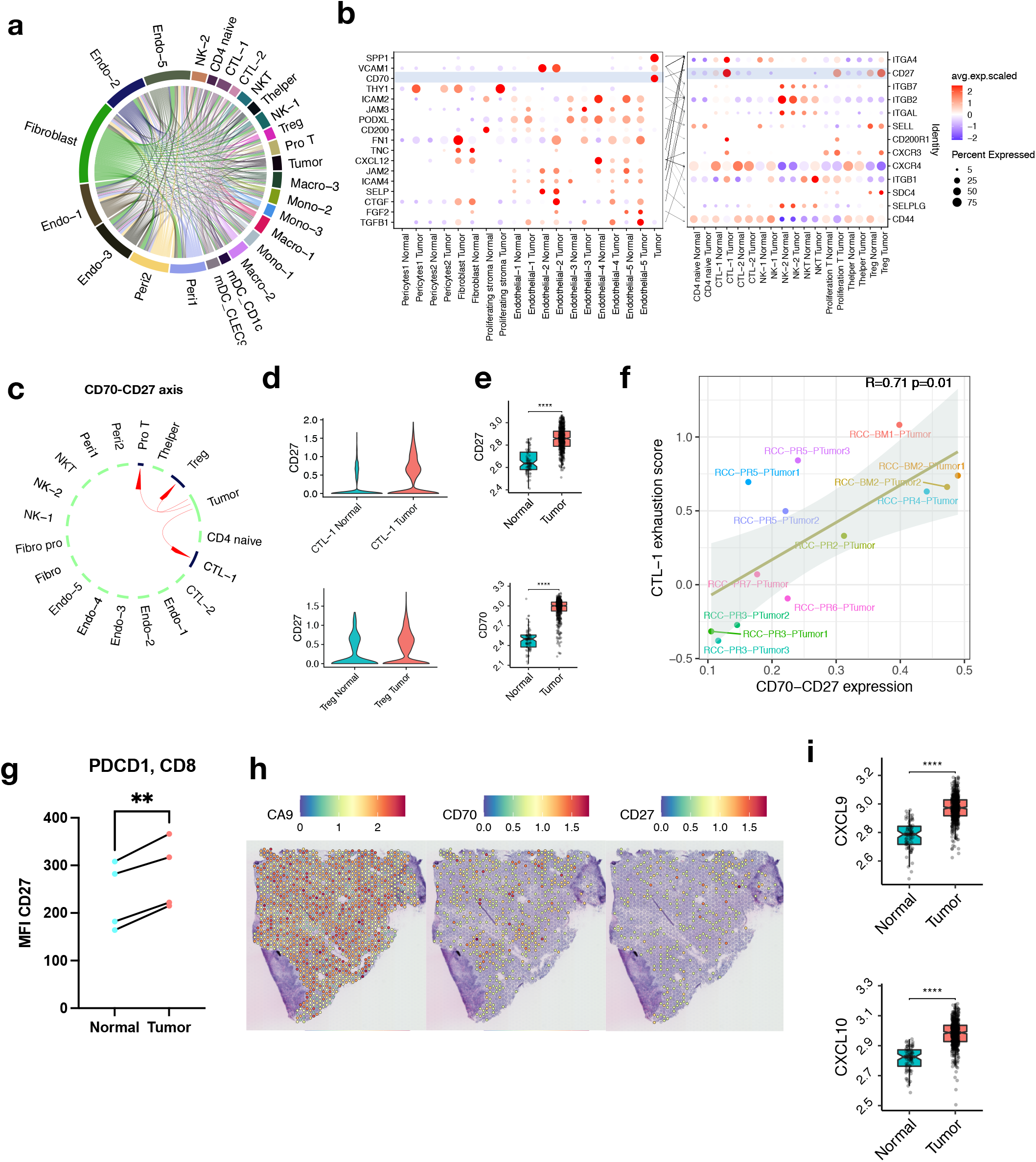
Cell-cell Interaction analysis reveals potential therapeutic targets in ccRCC. **a-** Overview of potential ligand-receptor interactions of cell subpopulations. **b**- Bubble heatmap showing expression of ligand (tumor cells and stromal cell subsets) and receptor (immune cell subsets) pairs in different stromal and immune subsets. Dot size indicates expression ratio, coloured represents average gene expression. Significance of ligand-receptor pair is determined by permutation test, gene differential expression analysis and specific cellular expression (Methods). **c**- The predicted interactions between CD70 and CD27. **d**- Violin plot showing CD27 expression in CTL-1 and Tregs. **e**- CD27 and CD70 expression in TCGA KIRC data are shown as boxplot. Statistics are accessed with Wilcoxon rank sum test. **f**- Correlation of CD70-CD27 (tumor cells-CTL-1) average expression and CTL-1 exhaustion score is shown as scatter plot. Pearson linear correlation estimate and p-values are shown. **g**- Flow cytometric analysis of CD27 expression on PDCD1+ CD8+ T cells from Tumor and paired adjacent normal tissue. (n= 4 per group). Statistics are accessed with paired t-test (** p < 0.001). **h**- Spatial feature plots showing CA9, CD70 and CD27 expression in ccRCC patient. Tumor spots are marked by CA9 expression. **i**- CXCL9 and CXCL10 expression in TCGA KIRC data are shown as boxplot. Statistics are accessed with Wilcoxon rank sum test (****p<0.0001). Boxplots include center line, median; box limits, upper and lower quartiles; and whiskers are highest and lowest values no greater than 1.5x interquartile range.

### ccRCC are highly enriched with vascular remodeling and EMT switch

Clear cell RCC are highly vascularized tumors with disorganized vasculature, endothelial cells and pericytes^59^. Subcluster analysis of the stromal compartment revealed two pericyte (*RGS5, ACTA2, TAGLN*), two fibroblast (*DCN, LUM*) and five endothelial (*CD34, VWF*) sub-populations (Fig. 4a, Extended data Fig. 6a, b). A notable proportion of the pericytes, and some of the endothelial and the fibroblast cells, were only detected in the tumor fraction (Fig. 4a, b), suggesting a tumor-induced remodeling of the stroma.

Pericyte-1 was enriched in the tumor fraction (Fig, 4e) and characterized by higher expression of genes related to a pro-inflammatory capillary phenotype (*RGS5, KCNJ8, AXL, ABCC9, PDGFRB, THY1*) (Extended data Fig. 6c). Capillary pericytes, named because they are wrapped around capillaries, favor vessel sprouting through the promotion of endothelial tip cell formation, stimulating tumor angiogenesis and possible hematogenous spread of tumor cells^60^. The Pericyte-2 cluster exhibited a vascular smooth muscle cell (VSMC) phenotype (*ACTA2, CNN1, RERGL MYH11, TAGLN, PLN*) ^61^ (Extended data Fig. 6c), and the proportion of cells was significantly reduced in the tumor compartment (Fig. 4e). To validate this, we analysis spatial transcriptomic data and found Pericyte-2 markers are less present in tumor infiltrated region compared to Pericyte-1 (Extended data 7a-c). VSMCs are thought to maintain the structural integrity of the blood vessels through their contractile properties and by inhibiting ECM degradation that would result from processes including vessel sprouting^61^. These pericyte changes may indicate that the vascular remodeling during RCC progression favors the pro-inflammatory and pro-invasive pericyte populations over the VSMCs^60,61^.

The fibroblast cluster was annotated based on the expression of marker genes such as *DCN* and *LUM* (Fig 4c; Extended data Fig. 6a). Compared to adjacent normal tissue, we found cancer associated fibroblast (CAF) signature genes^62^ (*FAP, FN1, LRRC15, THY1 and TGFBI*) were increased in the tumor (Fig 4h; Extended data Fig. 6e) and associated with poor prognosis in two independent RCC cohorts (Fig 4i). Interestingly, we detected a distinct stromal sub-cluster with a proliferating phenotype (*MKI67*), which was enriched in the tumor compartment (Fig. 4a-c, Extended data Fig. 5e). However, the data is limited to make further conclusions of this population.

The endothelial population appear to be significantly decreased in tumor compared to adjacent normal kidney tissue (Extended data 1c; Extended data 7d). Within the endothelial cells, endothelial-1 (Endo-1) was the most abundant and enriched in the tumor (Fig. 4a-b, 4f), and characterized by an upregulation of tumor-associated genes (*PVLAP, CA2, SPARC, INSR, IGFBP7*)^63^ (Fig. 4c, Extended data Fig. 6b, d). This gene signature is consistent with a tumor-associated endothelial cell (TEC) phenotype, which has been previously described and characterized by upregulated expression of pro-angiogenic factors^64^. Interestingly, a subcluster of Endo-1 was found almost exclusively in the tumor compartment (Fig. 4b) preferentially expressing known endothelial tip cell genes (*KCNE3, DLL4, EDNRB, ANGPT2, SERPINE1*) (Extended data Fig. 6b, f). These endothelial tip cells coordinate the sprouting of capillaries from pre-existent vessels and create new access points for tumor cells into the blood stream^63^. In a spatial context, we also observed high expression of *PLVAP* and *CA2* within tumor infiltrated region (Extended data 7b-c).

Endothelial-2 (Endo-2) cells expressed genes of the vascular endothelium (*DARC, VCAM1, VWF*) (Fig. 4c) and preferentially expressed venous EC genes (*GPM6A, CYP1B1, MMRN1*) (Extended data Fig. 6b, d)^63^. Moreover, upregulation of genes *CLU, NNMT* and *S100A6*, which are known to promote RCC cell proliferation and metastasis, were found exclusively in the tumor compartment of Endo-2 (Extended data Fig. 6b, d) and could indicate a supportive role of these cells to RCC progression^65,66^. Endothelial-3 (Endo-3) was characterized by high expression of *EHD3* (Fig. 4c) which is known to be expressed exclusively by kidney glomerular ECs (GECs)^67^. Interestingly, Endo-3 cells were abundant in normal adjacent tissue but significantly decreased in the tumor (Fig. 4f) and showed an increased angiogenesis score (Fig. 4d). GECs are crucial to the integrity of the glomerulus and damage to these cells contributes to the progression of chronic kidney disease^67^. This cluster showed high expression of the angiogenesis-related endothelial hematopoietic gene *EMCN* and *MEG3* (Fig. 4c), which has been shown to inhibit cell inflammation in RCC^68,69^.

The endothelial-4 (Endo-4) cluster was characterized by expression of hematopoietic stem cell-supporting factors (*CXCL12, CD44*). The endothelial-5 (Endo-5) cluster showed a tumor-associated EC phenotype (*IGFBP3, ENPP2, SEMA3G, TM4SF1, TIMP3*) (Fig. 4c; Extended data Fig. 6b, d), high angiogenesis score (Fig. 4d) as well as similar upregulation of pathways related to blood vessel and circulatory development (Fig. 4g)^63^.

### Ligand-receptor cell interaction analysis reveals potential novel therapeutic targets in human ccRCC

Although renal tumors are highly infiltrated with immune cells, RCC often successfully evades immune recognition by poorly understood mechanisms^13^. We interrogated ligand-receptor interactions with the goal of identifying new prognostic biomarkers and therapeutic targets (Fig. 5a). Several ligands were specifically upregulated in the tumor cell population when compared to adjacent kidney, including *SPP1* and *CD70* (Fig. 5b).

*SPP1*, also known as Osteopontin (OPN), promotes cancer progression and metastasis through activation of NFkB signaling which regulates extracellular matrix interactions^70^. Elevated levels of *SPP1* in ccRCC are correlated with larger tumor size, advanced stage, higher Ki-67 proliferation index, and decreased overall survival^70^. *SPP1* binds immune cells including macrophages, NK cells and T cells, and exerts immunomodulatory actions^71^. The *SPP1* receptor *ITGB1* was upregulated in tumor derived NK cells while another *SPP1* receptor *ITGA4* was upregulated in tumor derived CTL-1 cells. This suggests that SPP1 produced by tumor cells exerts tumor-mediated immunoregulation through NK cells and T cells (Fig. 5b).

The ligand *CD70* induces apoptosis in B and T lymphocyte populations in ccRCC and has been associated with immune suppression^72^. We found that CD70-CD27 signaling was upregulated in tumor tissue compared to adjacent kidney (Fig 5c, e; Extended data Fig. 8b, d-g). CD70-CD27 expression correlated with T cell exhaustion (Fig. 5f). The *CD27* receptor was overexpressed in tumor-associated CTL-1 (Fig. 5b-d, g; Extended data Fig. 8a, c), and correlated with CTL-1 exhaustion (Extended data 9a, c). Both the ligand, CD70, and its receptor, CD27, are independently expressed in tumor tissue (Fig. 5h; Extended data 9d). Our data suggests that CD70 is immunosuppressive and that its receptor CD27 merits exploration as a therapeutic target in ccRCC^73^.

The chemokines CXCL9 and CXCL10 and their receptor *CXCR3* (expressed on monocytes, T and NK cells) appear to be involved in angiogenesis^74^. High expression of *CXCR3* and *CXCL9/10* has been associated with a poor prognosis, tumor growth, and increased risk of metastasis^75^. Our analysis suggested that *CXCL9-*expressing mDC_LAMP3 cells, as well as *CXCL10*-expressing Macro cells, interact with CXCR3-expressing proliferative T cells. In addition, *CXCL9*- and *CXCL10*-expressing Macro-2 tumor cells interact with *CXCR3*-expressing proliferating T cells, T_regs_ and CTL-1 cells in the tumor (Fig. 5i; Extended data 8h; Extended data 9e, g). We demonstrated overexpression of *CXCL9* in mDC_LAMP3 and Macro-2 cells, and overexpression of *CXCL10* in proliferating macrophages (Pro Macro) and Macro-2 cells in the tumor (Extended data 8h; Extended data 9f). We observed that *CXCL9 and CXCL10* are correlated with CTL infiltration and PDCD1 expression in multiple clinic cohorts from TIDE^76^ (Extended data 9h). We therefore hypothesize that *CXCL9/10* signaling via *CXCR3* may impact the microenvironment and potentially promote tumor progression through deregulation of inflammatory pathways.

## DISCUSSION

Our study took advantage of coordinated surgical and research teams to bring fresh patient samples from the operating room to the laboratory. This enabled the single-cell characterization of treatment naïve ccRCC primary tumors and adjacent normal kidney tissue with a focus on the tumor microenvironment. We uncovered intratumor heterogeneity, as well as a highly heterogenous tumor microenvironment, and multiple immunosuppressive cell-cell interactions. Further investigation of these cell-cell interactions in the tumor ecosystem highlighted novel potential therapeutic targets.

Clear cell RCC tumor cells almost universally display the loss of function of the von Hippel-Lindau (VHL) gene. Consistent with this, we observed that malignant cells have recurrent deletions of chr3p (where *VHL* is located) and an upregulation of *VEGFA* and *VIM*. The cellular origin of ccRCC has been suggested to be from proximal tubule epithelium^3^. Indeed, malignant cells show similar transcription profiles with the PT2 cluster (Fig. 3a) pointing towards the PT2 subset as the ccRCC cell of origin. Intratumor heterogeneity has been widely reported^77,78^ and we identified 4 subsets of malignant cells. Notably, malignant cell cluster 4 was associated with poor overall survival and was enriched in patients with metastases. Cluster 4 was characterized by expression of *SAA1, SAA2* and *APOL1*, genes that are upregulated in high grade RCC tumors. Indeed, high *SAA1* protein expression has been negatively correlated with patient survival^58^. We suggest that Cluster 4 cells exhibit a distinct transcriptional signature that portends metastatic potential, and a specific focus on this cluster may highlight new therapeutic targets to prevent or treat metastatic spread.

Within the complex immune microenvironment, we identified exhausted T cells (CTL-1) and immunosuppressive cell populations including T_regs_ and Macro-2. During cancer progression, CTLs can exhibit loss of function as they become exhausted due to immune related tolerance and immunosuppression. Cancer-associated fibroblasts (CAFs), M2-macrophages and T_regs_ may counteract the CD8^+^ T and NK cell mediated antitumor immune responses^79^. Studies in melanoma have shown that increased expression of *PD-L1, IDO* and *FoxP3* protein in T_reg_ of the TME is driven by infiltration of CD8^+^ T cells, further supporting the idea that these are part of an immune negative feedback loop^41^. Immunotherapies that uncouple these pathways may be the most effective on tumors showing T cell infiltrated phenotypes. In RCC, T cell infiltration into the TME has been demonstrated^11,12^ and anti-*CTLA-4* and anti-*PD-1* antibodies are approved treatment strategies in patients with advanced disease^15,80^. Both *PD-1* and *CTLA-4* receptors are upregulated in T_regs_ and contribute to the immunosuppressive function of Tregs^81^. T cell immunoglobulin mucin-3 (*TIM-3/HAVCR2*) is another immune checkpoint surface receptor present on T_regs_ that tends to suppress the immune responses^82^ and therefore resist radiation therapy^83^. This particular immunosuppressive cell population of T_regs_ may represent another potential therapeutic target in ccRCC.

Regarding myeloid cells within the TME, blocking the polarization to M2 macrophages results in increased recruitment of CTLs and an antitumor immune attack^84^. Conversely, induction of M2 macrophages impairs the response of CTLs in the TME^84^. We show that this phenomenon also exists in ccRCC, as illustrated by the increased Macro-2 (Fig. 2b) and the increased exhaustion score of the CTL-1 (Fig. 2f). The Macro-2 population expressed high levels of *TREM2* in the tumor fractions and expression of *TREM2* has been shown to be elevated in malignant tumors, including RCC^85^. *TREM2*^*-/-*^ mice are more responsive to immune checkpoint blockade and less susceptible to cancer progression^36^. *TREM2* inhibition of the tumor-infiltrating macrophages suppressed tumor growth and enhanced checkpoint blocking therapy in preclinical studies. Here, we show that *TREM2* is elevated in the TME myeloid cells (Extended data 2g-i), pointing to the potential as a prognostic marker in ccRCC and as a biomarker that could identify patients that would benefit from checkpoint blockade immunotherapy. Our interactome analysis suggests that the immunosuppressive role of *TREM2* macrophages (Macro-2) affects T cell exhaustion through C-X-C motif chemokine ligand 9/10 (*CXCL9/CXCL10*) and CXC-chemokine receptor 3 (*CXCR3*) signalling^36,86^.

In terms of cell-cell interactions, we identified the CD70-CD27 relationship as potentially important, hypothesizing that up-regulated CD70-CD27 signaling in the tumor may result in CTL exhaustion. Here, the tumor cells expressed the ligands *CD70* and *CD80–CD86* that bind to their respective *CD27* and *CD28* receptors expressed on CD8^+^ T cells. The receptor-ligand interactions represent the first step for CD8^+^ T cell priming^87^. When activated, naive CD4^+^ and CD8^+^ T cells upregulate *CD27*, resulting in increased circulating soluble *CD27*, a diagnostic marker of T-cell activation^88^. CD70, the ligand of *CD27*, is exclusively expressed after immune activation and is also found in both CD8^+^ T cells and T_regs_^88^. The ligand CD70 is frequently overexpressed in several solid cancers, most prominently renal cancer (87%) but also including lung cancer (10%), glioblastoma (42%), and ovarian cancer (15%). Given that its receptor CD27 is predominantly found on exhausted T-cells (CTL-1) (Fig. 5g; Extended data 8e-g; Extended data 9a, c-d), CD70-CD27 interactions may contribute to immune evasion by tumor. If so, these interactions, would represent a potential therapeutic target^89^.

In conclusion, our study provides several important biological insights about RCC: (1) We identify a distinct metastatic signature that predicts survival outcome, ^90^ we characterized a proximal tubule cell population that may represent the RCC cell of origin, and (3) we dissect the immunosuppressive environment and the stromal alterations in treatment-naïve patients. We highlight potential therapeutic targets to be further evaluated in preclinical models. We hope that our study will provide a valuable resource for further explorations into critical cellular relationships and signaling axes that remodel the tumor microenvironment resulting in a tumor-suppressive environment. Ultimately, validating these relationships may lead to new approaches to address the clinical problem of relapsed and metastatic RCC.

## Acknowledgments

We are particularly indebted to our patients and their clinical care teams. We gratefully acknowledge support from Bill & Cheryl Swanson, Gunther & Maggie Buerman, and Robert Higginbotham. A.A. and N.B. was funded by the Swedish Cancer Society. P.V.K. was funded by the NIH grant R01HL131768 from NHLBI. Patient samples were sorted at the HSCI/CRM flow cytometry core facility at MGH with the help of Maris Handley and Pathik Sen.

## Author contributions

Conceptualization, A.A., S.M., P.S., D.B.S., P.V.K., and N.B.; Investigation, A.A., S.M., I.T., T.H., Y.K., P.S., D.B.S., C.W., A.S., S.W., P.V.K., and N.B.; Computational investigation and analysis, A.A., S.M., I.T., P.V.K., and N.B.; Writing -– Original draft, A.A., S.M., I.T., and N.B.; Writing – Review & Editing, A.A., S.M., C.W., A.S., S.W., P.S., D.B.S., P.V.K., and N.B.; Funding Acquisition, Resources & Supervision, P.S., D.B.S., P.V.K., and N.B.

